# CD13 is a Critical Regulator of Cell-cell Fusion in Osteoclastogenesis

**DOI:** 10.1101/2020.04.25.061325

**Authors:** Mallika Ghosh, Ivo Kalajzic, Hector Leonardo Aguila, Linda H Shapiro

## Abstract

In vertebrates, bone formation is dynamically controlled by the activity of two specialized cell types: the bone-generating osteoblasts and bone-degrading osteoclasts. Osteoblasts produce the soluble receptor activator of NF*κ*B ligand (RANKL) that binds to its receptor RANK on the surface of osteoclast precursor cells to promote osteoclastogenesis, a process that involves cell-cell fusion and assembly of molecular machinery to ultimately degrade the bone. CD13 is a transmembrane aminopeptidase that is highly expressed in cells of myeloid lineage has been shown to regulate dynamin-dependent receptor endocytosis and recycling and is a necessary component of actin cytoskeletal organization. In the present study, we show that CD13-deficient mice display a normal distribution of osteoclast progenitor populations in the bone marrow, but present a low bone density phenotype. Further, the endosteal bone formation rate is similar between genotypes, indicating a defect in osteoclast-specific function in vivo. Loss of CD13 led to exaggerated in vitro osteoclastogenesis as indicated by significantly enhanced fusion of bone marrow-derived multinucleated osteoclasts in the presence of M-CSF and RANKL, resulting in abnormally large cells with remarkably high numbers of nuclei with a concomitant increase in bone resorption activity. Similarly, we also observed increased formation of multinucleated giant cells (MGC) in CD13^KO^ bone marrow progenitor cells stimulated with IL-4 and IL-13, suggesting that CD13 may regulate cell-cell fusion events via a common pathway, independent of RANKL signaling. Mechanistically, while expression levels of the fusion-regulatory proteins dynamin and DC-STAMP are normally downregulated as fusion progresses in fusion-competent mononucleated progenitor cells, in the absence of CD13 they are uniformly sustained at high levels, even in mature multi-nucleated osteoclasts. Taken together, we conclude that CD13 may regulate cell-cell fusion by controlling expression and localization of key fusion proteins that are critical for both osteoclast and MGC fusion.

## Introduction

Osteoclastogenesis is a critical process for skeletal growth and development that is tightly regulated by differentiation of myeloid progenitor cells into osteoclasts, which are specialized cells in the bone marrow whose major function is bone resorption^1^. This highly-regulated process is responsible for bone modeling and remodeling that ultimately translates into maintenance of bone integrity, skeletal growth and repair^2,3^. Deregulation of the process leads to dramatic outcomes^4,5^. Loss of osteoclast (OC) generation/function gives rise to elevated bone formation without remodeling, resulting in osteopetrosis characterized by high bone mass and growth impairment, while gain of OC generation/function results in exacerbated bone degradation that, without equivalent coupling to bone formation, leads to osteoporosis, characterized by low bone mass, bone weakness and high predisposition to fractures with poor healing progression. OCs form from committed monocyte progenitors via initial signals provided by two central bone marrow-derived cytokines: macrophage colony stimulating factor (M-CSF) and receptor activator of NF*κ*B ligand (RANKL), initiating a highly-organized program of commitment towards terminal differentiation and function^6-8^. This process involves proteins that participate in cell-cell fusion which are critical to the generation of functionally active OCs. Despite our current knowledge of events leading to the formation of OCs, understanding the regulation of these processes in homeostatic as well as their de-regulation in pathological conditions is unclear^9-11^.

CD13 is a transmembrane metalloprotease widely expressed in all cells of the myeloid lineage, as well as activated endothelial cells, hematopoietic progenitor and stem cells^12-16^. Previous studies have implicated CD13 in vasculogenesis, tumor cell invasion, inflammatory trafficking and as a receptor for human corona virus^17-19^. Our novel observations have clearly shown that independent of its peptidase activity, CD13 is a critical molecule that assembles the molecular machinery enabling diverse cellular processes such as cell-cell adhesion, migration, membrane organization and dynamin-mediated receptor endocytosis and recycling of cell surface proteins^14,15,20-22^. Taken together, CD13 regulates many of the activities that have been described to be critical for osteoclastogenesis and cell-cell fusion. In the current study, we demonstrate that despite a relatively normal distribution of hematopoietic components in bone marrow and periphery, CD13^KO^ mice have reduced bone mass with increased OC numbers per bone surface area but normal bone formation parameters, indicating that osteoclastogenesis is compromised in the absence of CD13. In addition, the in vitro induction of CD13-deficient myeloid progenitors generated from bone marrow and spleen resulted in increased numbers of OCs that were considerably larger in size, contained many more nuclei and resorbed bone more efficiently than those from wild type progenitors. We confirmed that CD13-deficient macrophages can also hyper- fuse to generate elevated multinucleated giant cells (MGCs) which again, were larger and contained more nuclei than those generated from wild type macrophages, suggesting that CD13 is a component of common fusion pathways shared by OC and MGC. Furthermore, we demonstrated that expression of fusion proteins which is typically downregulated in mature osteoclasts post-fusion^23^ is abnormally preserved in osteoclasts lacking CD13. These findings are in agreement with our data showing that CD13 is a mediator of homotypic cell interaction and a regulator of molecular events defining cell membrane organization, fluidity and movement, all processes critical to cell-cell fusion^14,15,20^. We hypothesize that CD13 is a negative regulator of cell-cell fusion in osteoclastogenesis and giant cell formation and potentially, a universal modulator of membrane fusion and is a novel target for therapeutic intervention in pathological conditions mediated by defects in cell-cell fusion.

## Results

### Bone density is reduced CD13^**KO**^ **mice *in vivo***

Based on the notion that the high, sustained expression of CD13 in all cells of the myeloid lineage reflects its important role in myeloid cell biology, we and others have demonstrated that it contributes to many fundamental cellular processes that impact myeloid cell function in various tissues. These studies prompted our current focus on the myeloid cells of the bone, the osteoclasts. We initially examined the effect of a global loss of CD13 on the phenotype of developing bone. Analysis of bone micro-architecture and function in the cortical and trabecular bone isolated from 8-10 wks. old WT and CD13^KO^ male mice by μ-CT and histomorphometric analysis revealed that the femur cortical and trabecular bone density and thickness in CD13^KO^ mice is significantly reduced (1.5 fold) compared to WT animals (**FIG. 1A, B**). **Immunohistochemical analysis of TRAP stained frozen sections of the femur clearly indicated elevated TRAP**^**+**^ **osteoclasts (in purple) in CD13**^**KO**^ **femur compared to the wildtype counterpart** (**FIG. 1C**). Reduced bone volume**/**total volume (BV/TV, **1D)**, number of osteoclasts per bone surface (OC/BS, **1E)** and per bone perimeter (OC/B.Pm, **1F)** with similar endosteal bone formation rates per bone surface (BFR/BS, **1G**), together, strongly suggested a defect in bone remodeling in the absence of CD13. Osteoblast cultures from BM stained for alkaline phosphatase activity indicated that alkaline phosphatase (ALP)^+^ osteoblast surface per well was equivalent between genotypes (data not shown) supporting a contribution of CD13 to osteoclast activity in vivo and in vitro.

**FIG 1.**
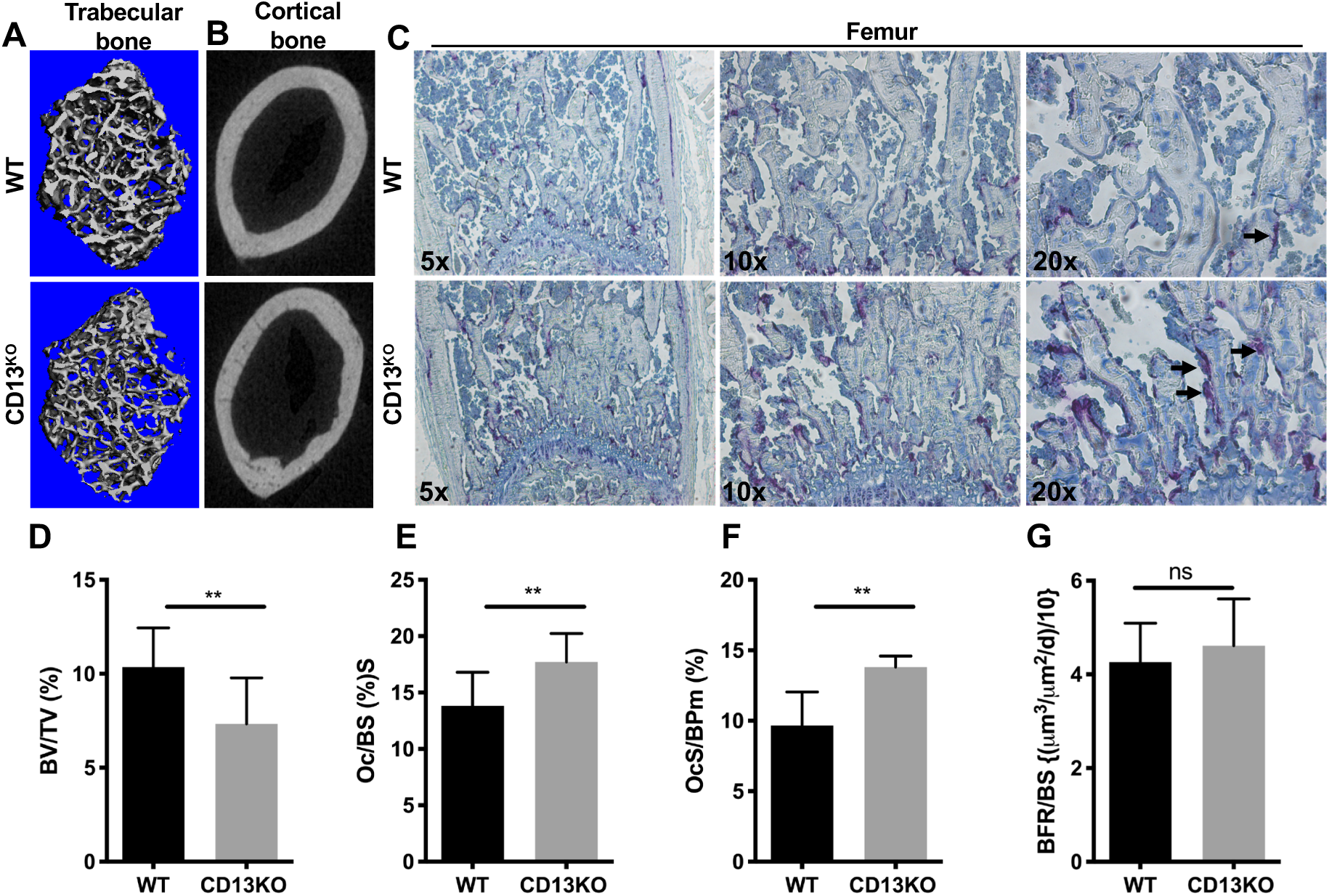
Loss of CD13 leads to reduced bone volume and increased number of osteoclasts in vivo. μCT and histomorphometric analysis of trabecular (**A**) and cortical (**B**) bone with (**C**), immunohistochemistry of TRAP^+^ osteoclasts in the femur indicated in purple (arrow), (**D**), BV/TV; Bone volume/Tissue volume (**E**), Oc/BS; % OCs per bone surface (**F**), OcS/BPm; % Osteoclasts per bone perimeter and (**G**), endosteal bone formation rate relative to bone surface area (BFR/BS) of WT and CD13^KO^ are shown. Data represents +/- SEM of two independent experiments. N=6/genotype. *\*\*;p<0*.*01*.

### OCP populations with osteoclastogenic potential are similar in WT and CD13^**KO**^ bone marrow

Previously we have shown that the distribution of the hematopoietic population comprised of early hematopoietic progenitors, myelo-erythroid progenitors, common myeloid progenitors, and granulocyte macrophage progenitors in CD13^KO^ mice were similar to wildtype animals^12^. Cells that can generate bone-resorbing osteoclast reside in both bone marrow and peripheral hematopoietic organs^24^. To determine if differences in osteoclast progenitor (OCP) frequency are responsible for the loss of bone mass in the absence of CD13, we analyzed the distribution of primary OCP in both the bone marrow microenvironment and spleen in WT and CD13^KO^ mice. Flow cytometric analysis (**FIG. 2A,B)** revealed that the OCP profile indicated by CD3-, B220-, NK1.1-, CD11b-/lo, CD115+, CD117+ in the BM (**A**, WT vs. CD13^KO^; 1.7 vs. 1.95) and CD3-, B220-, NK1.1-, CD11b+, Ly6G-, Ly6C+, CD115+ in spleen (**B**, WT vs. CD13^KO^; 0.24 vs. 0.238) ^24^ is similar between genotypes, indicating that the absence of CD13 does not change the differentiation potential of myeloid cells to osteoclast progenitors.

**FIG 2.**
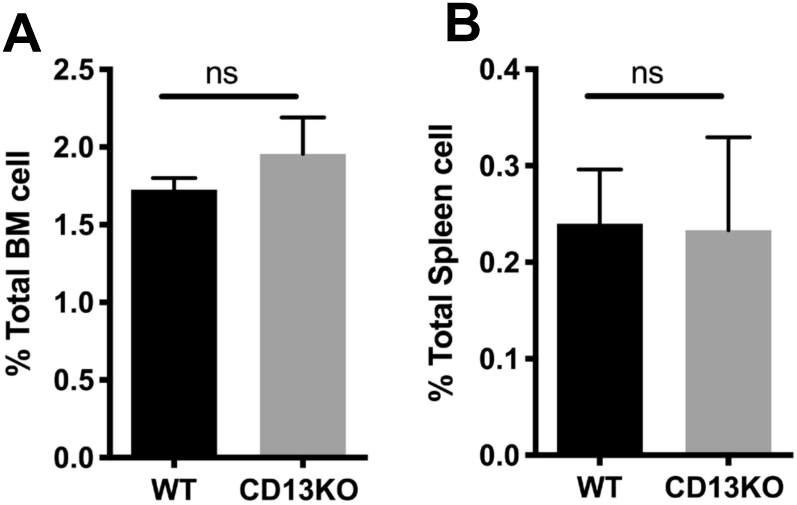
Differentiation of hematopoietic cells to osteoclast progenitors is independent of CD13. Flow cytometric analysis of Osteoclast progenitor population from BM (**A**) and Spleen (**B**) indicate similar distribution of progenitor cells between WT and CD13^KO^ mice. Data represents +/- SEM of three independent experiments. N=6/genotype.

### In vitro osteoclastogenesis is exaggerated in CD13^**KO**^ cells

Stimulation of monocyte lineage-committed hematopoietic progenitors with two principal bone marrow cytokines, M-CSF and RANKL, triggers the expression of molecules involved in cell-cell fusion and functional bone resorption^6-8^ and the generation of multinucleated osteoclasts. The normal hematopoietic profiles in CD13^KO^ mice, taken together with the defective skeletal phenotype, suggested that CD13 may participate in osteoclastogenesis in vitro. Analysis of flow sorted, bone marrow-derived OCP (CD3- B220- NK1.1- CD11b -/lo CD115+ Ly6C+, **FIG. S1A**) differentiated to mature OC in the presence of recombinant M-CSF (30ng/ml) and RANKL (30ng/ml) stained with TRAP, revealed significant increases in the number of osteoclasts containing >3 nuclei per field (2.8 fold), the average OC size (4-fold) and the number of nuclei/OC (2.5-fold) in CD13^KO^ cells grown on plastic (not shown) or bovine cortical bone slices (**FIG. 3A-D**) compared to those generated from normal WT progenitors. This difference was evident by d3, suggesting that the lack of CD13 accelerates OC fusion and multinucleation but not OCP proliferation rates, as the cell density (total number of nuclei/dish) was not significantly different between genotypes (**FIG. 3G**). Furthermore, we examined the flow sorted spleen-derived OCP (CD3- B220- NK1.1- CD11b+ Ly6G- Ly6C+ CD115+), which were allowed to proliferate in presence of M-CSF for d3, followed by differentiation into mature OCs by RANKL treatment for an additional 3 days. Similar to BM-derived cells, CD13^KO^-derived splenic cells produced OCs with larger area (3-fold), more cells with >3 nuclei (2.5-fold) and an increased number of nuclei per cell (3-fold) compared to WT mice (**FIG. 4A-D**). To assess OC bone resorptive capacity, we plated WT and CD13^KO^ flow-sorted bone marrow-derived OCP on Osteo assay plates (Corning) in the presence of recombinant M-CSF and RANKL and allowed them to mature to OC over time. At d10, OCs were removed, individual or multiple resorption pit areas were imaged and the area of resorption quantified by ImageJ. As expected, the increase in OC nuclei/cell positively correlates with resorption and CD13^KO^ OCs showed increased resorption areas (3-fold) compared to WT, confirming that the elevated osteoclastogenesis in CD13^KO^ mice translates into exaggerated functional activity (**FIG. 3E, F**). Our in vitro data is consistent with our in vivo bone phenotype in the absence of CD13.

**FIG 3.**
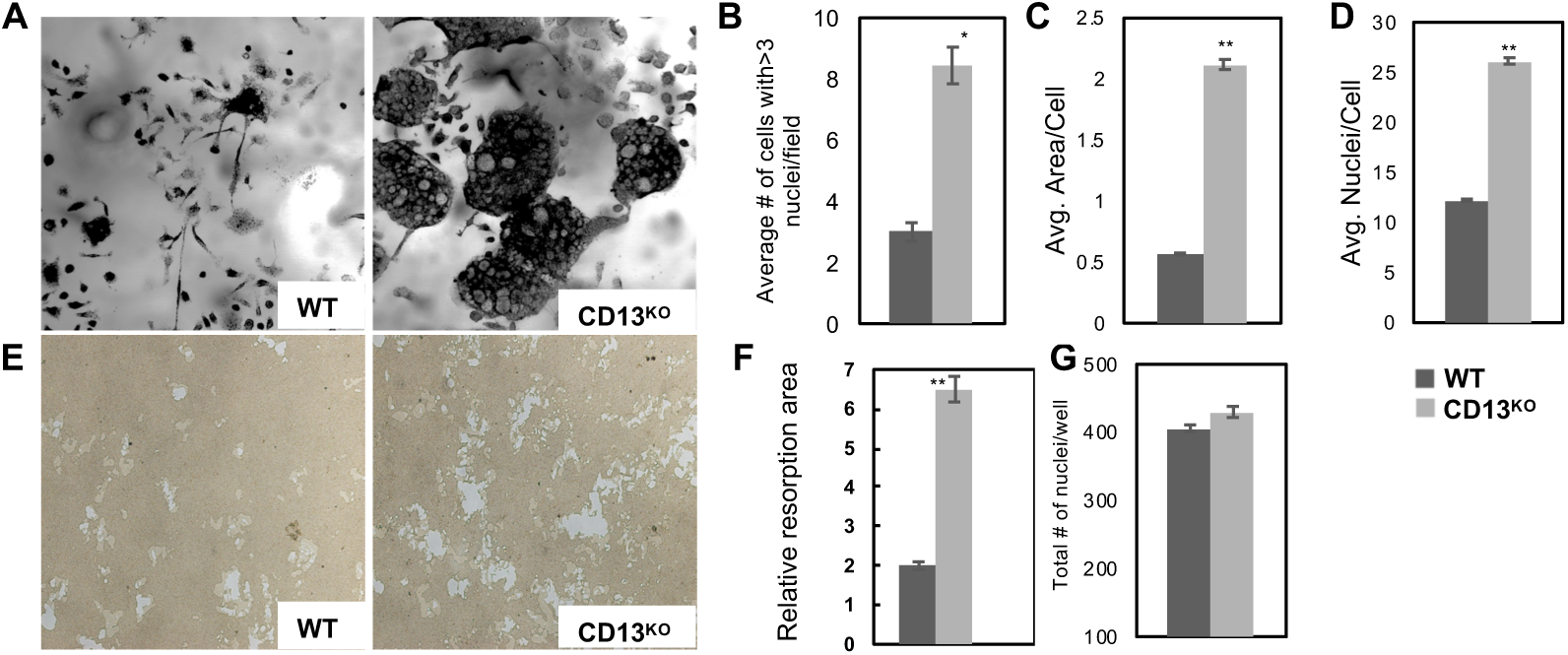
Increased TRAP+ multinucleated OCs in absence of CD13 in vitro. **A)**. Primary murine bone marrow derived OCs size and number of nuclei/OC grown on bovine cortical bone slices in CD13^KO^ cells compared to WT at d3. **B**). Average # of cell with> 3 nuclei/field, **C**). Average cell area and **D**). average size of nuclei per OCs in CD13^KO^ are significantly larger than the WT cells. **E**). Area of resorption is significantly higher in CD13^KO^ than WT OCs grown on Osteo assay plates (Corning) for d10 by phase contrast imaging & quantified by Image J (**F**). Data represents +/- SEM of three independent experiments. N=6/genotype, **;p<0.01, *;p<0.05.

**FIG 4.**
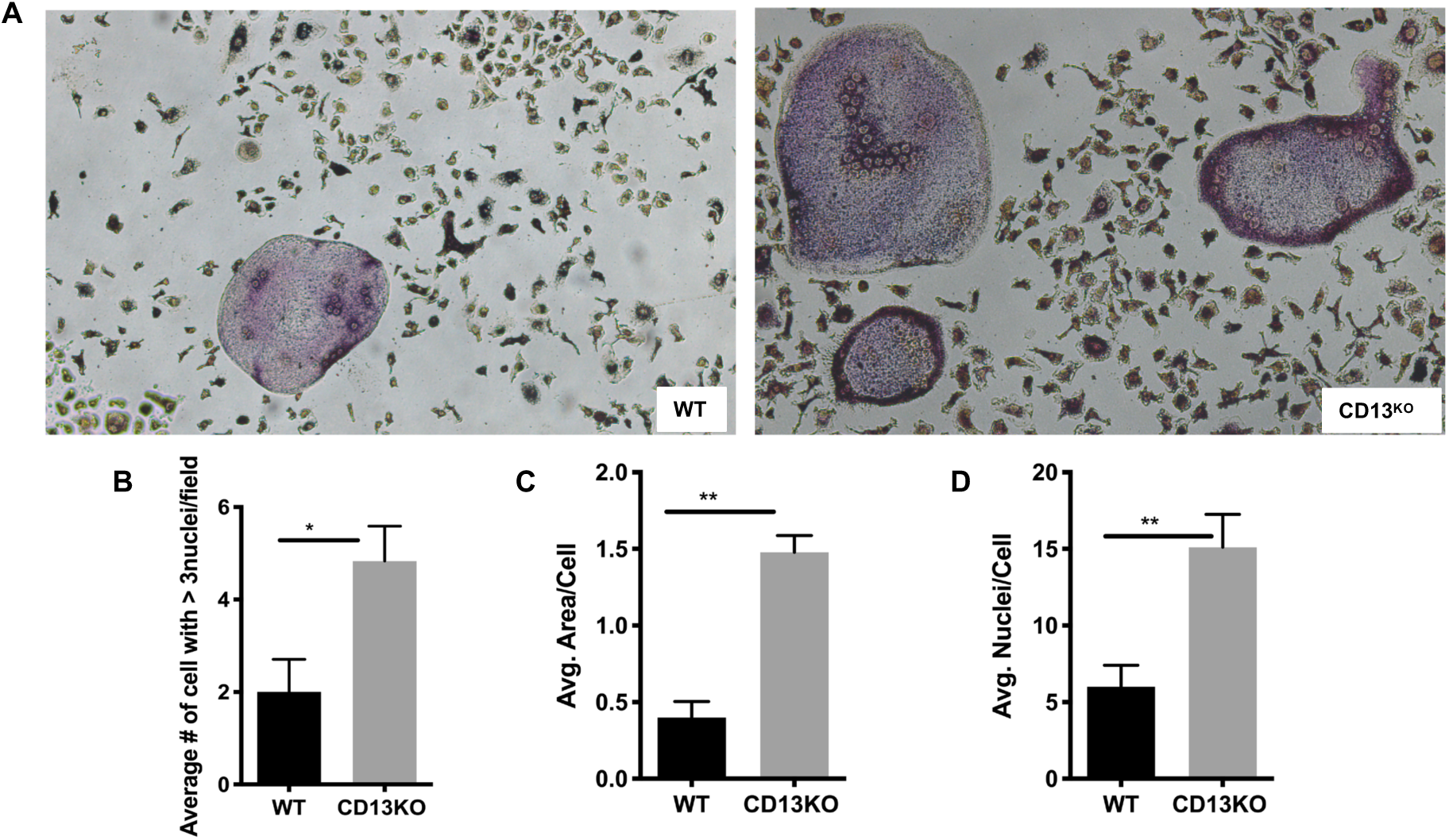
Increased TRAP+ spleen derived multinucleated OCs in absence of CD13 in vitro. **A)**. Primary murine spleen derived OCs size and number of nuclei/OC grown on dentine slices in CD13^KO^ cells in presence of M-CSF and RANKL compared to WT at d3. **B**). Average # of cell with >3 nuclei/field, **C**). cell area and **D**) average size of nuclei per OCs in CD13^KO^ are significantly larger than the WT cells. Data represents +/- SEM of two independent experiments. N=6/genotype, **;p<0.01, *;p<0.05.

### Macrophage giant cell formation (MGC) is enhanced in CD13^**KO**^ cells

In addition to osteoclasts and independent of RANKL, monocyte progenitors can differentiate into macrophages which also undergo cell-cell fusion to form large, multi-nucleated macrophage giant cells (MGC)^25,26^. To evaluate if CD13 also regulates MGC fusion, we expanded sorted BM myeloid progenitor cells (CD3- B220- NK1.1- CD11b -/lo CD115+ Ly6C+) in M-CSF for 5 days, followed by the addition of IL-4 (30ng/ml) and IL-13 (30ng/ml) for 3 days to promote macrophage fusion. Images of Giemsa stained cells indicated a significant increase in the number of cells per field with >3 nuclei (2.5-fold), increased area/cell and a greater number of nuclei/cell (5-fold) of MGCs in CD13^KO^ cultures compared to WT (**FIG. 5A-D**). Immunofluorescence and confocal microscopy further confirmed CD13 expression in CD68+ MGC (**FIG. 5E**). In agreement with CD13 as a cell surface mediator of fusion in macrophages, CD13 and the fusion protein DC-STAMP (DCST1)^27^ are co-localized in MGC stimulated with IL-4/IL-13 (**FIG. 5F**). Our data strongly suggest that CD13 regulates a common cell fusion event independent of RANKL signaling^28^.

**FIG 5.**
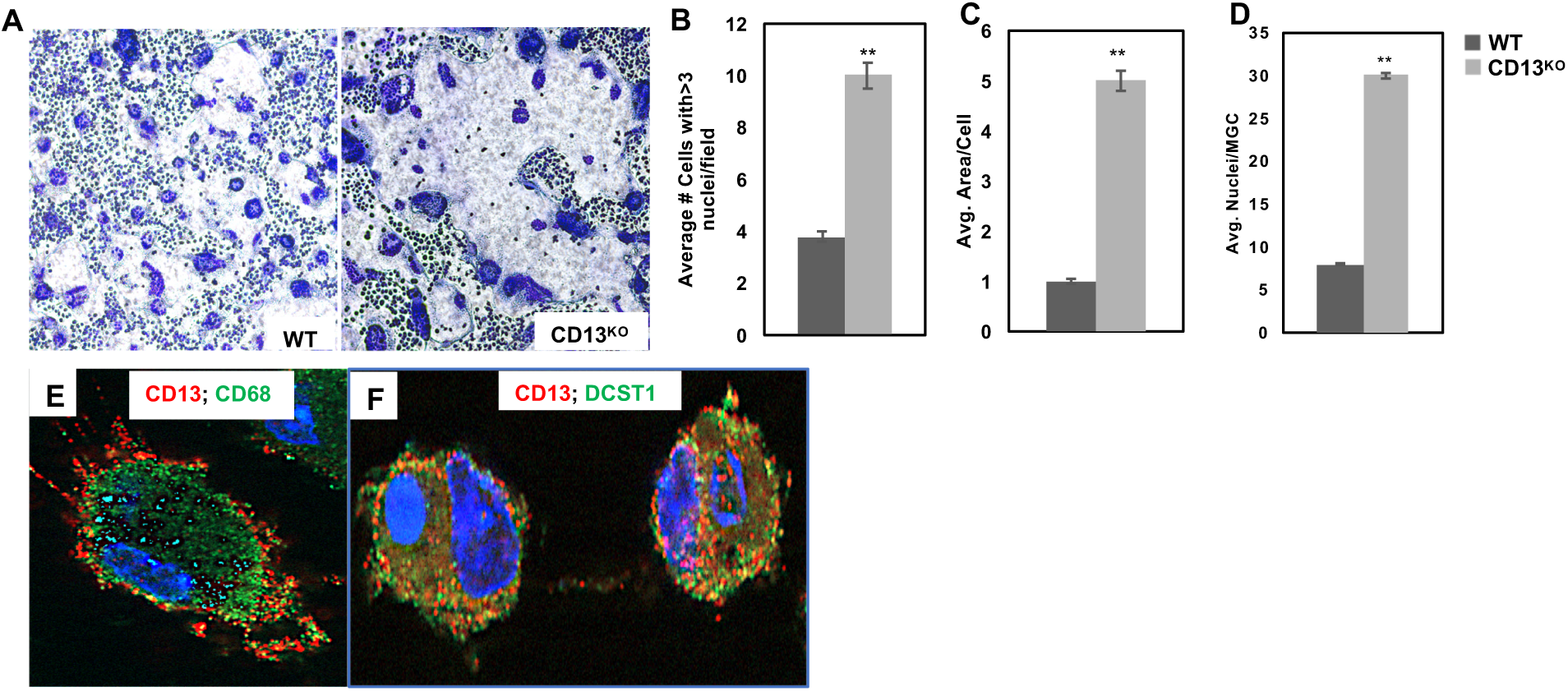
Increased Multinucleated Giant Cell formation in CD13^**KO**.^ **A**). Giemsa staining indicated increased size and number of MGCs in CD13^KO^ compared to wildtype counterpart in presence of M-CSF and IL-4/IL-13 after d5. **B**). Quantification of MGC with > 3 nuclei/field **C**). Average area and **D**). Number of nuclei/MGC. Data represents +/- SEM of two independent experiments. N=6/genotype, **;p<0.01. **E**). CD13 is expressed in CD68^+^ WT MGC and **F**). co-localize with fusion protein DCST1 after IL-4/IL-13 addition. CD68; green. CD13; red. Magnification 63X oil.

### Expression of fusion proteins is dysregulated in CD13^**KO**^ mature osteoclasts

Pertinent to our observations, it is believed that since OCs and macrophages are derived from common progenitors, some of the molecular mechanisms mediating their fusion and multinucleation may be shared^28^. In particular, the small GTPase dynamin 2 (the major isoform in OC) and the fusion protein DC-STAMP are common and critical regulators of both osteoclast and MGC fusion^23,27^. Our recent studies have shown that CD13 is a potent negative regulator of dynamin-dependent endocytosis of a variety of receptors^20-22^, suggesting that CD13 may participate in cell fusion by regulating endocytic processes. Indeed, immunoblot (**FIG. 6A**) and immunofluorescence (**FIG. 7**) analyses of flow-sorted WT BM-OCPs differentiated to OC with M-CSF and RANKL demonstrated high levels of dynamin and DC-STAMP expression by d2-post differentiation, which was subsequently reduced by 3d, when cell fusion and maturation into multinucleated WT osteoclasts is complete, as previously reported^23^. However, while dynamin and DC-STAMP are highly expressed in CD13^KO^ OCP, rather than being downregulated, this strong expression is maintained in mature multinucleated OCs (**FIG. 6A,7**), suggesting that CD13 may impact fusion by regulating the levels of key fusion and/or endocytic molecules critical for OC and MGC fusion. In addition, immunoblot analysis of cell lysates obtained from WT bone marrow-derived progenitor cells stimulated with M-CSF and RANKL over 3d indicated that CD13 is highly expressed in myeloid progenitor cells but its expression level is unaltered upon stimulation with M-CSF and RANKL over time (d0-3) (**FIG. 6B**), consistent with CD13 regulating fusion mechanisms independent of RANKL signaling. Further investigation into the relationship between CD13 and these and other common regulators of myeloid cell fusion such as OC- STAMP, the macrophage fusion receptor, osteoclast receptor αvβ3 integrin^28^ will clarify common mechanisms regulating CD13-dependent myeloid cell fusion as well as the fusion of other lineages including satellite stem cells.

**FIG 6.**
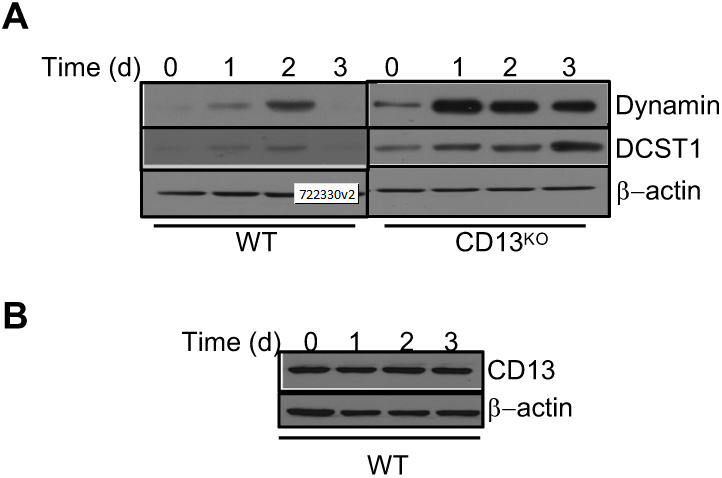
Evaluation of CD13 and fusion-promoting protein expression in M-CSF/RANKL stimulated BM-derived osteoclast by Immunoblot analysis. (**A**). Expression of the fusion- promoting proteins dynamin and DC-STAMP (DCST1) is aberrantly sustained in CD13^KO^ but not in WT multinucleated osteoclasts in presence of M-CSF and RANKL. (**B**). CD13 expression is unaltered in BM-derived cells in response to M-CSF and RANKL stimulation over time. Data represents average of two isolates. N=2/genotype.

**FIG 7.**
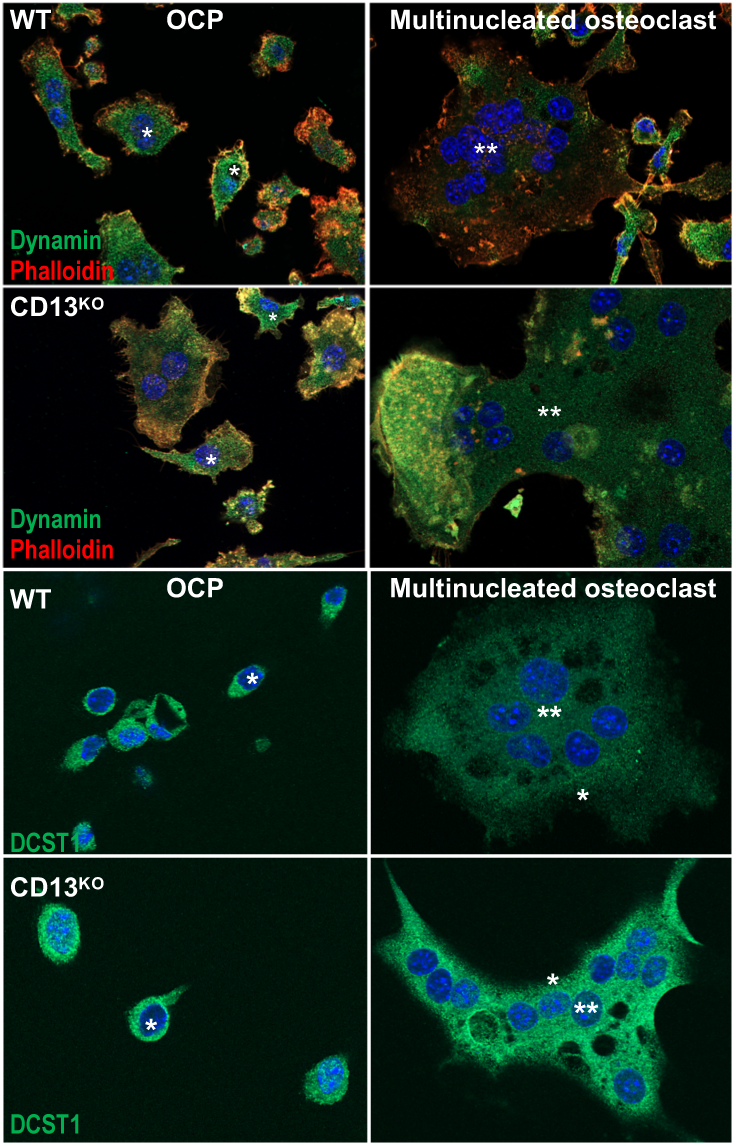
Immunofluorescence analysis of Dynamin and DC-STAMP (DCST1) in OC. Fusion proteins dynamin and DCST1 expression is maintained in CD13^KO^ multinucleated mature osteoclasts but not in WT cells (**). High level of dynamin colocalizes with actin and DCST1 in OCPs (*). Data represents average of three independent experiments. N=3/genotype. Dynamin, DCST1;green. Phalloidin; red. Magnification 63X oil.

## Discussion

The fusion of plasma membranes is essential to and indispensable for many physiologic processes such as fertilization through sperm/egg fusion^29^, muscular development through myoblast fusion^30^, skeletal development and maintenance of skeletal integrity through formation of multinucleated osteoclasts (OC); and control of certain viral infections and spreading through the formation of macrophage multinucleated giant cells (MGC)^31^. Thus, this important biological process directly defines the course of many pathological processes including infertility, skeletal defects (osteoporosis and osteopetrosis), failure of skeletal repair, failure of maintenance of prosthetic implants as well as fusion of host plasma membrane and virus fusion protein in viral diseases. Clearly, potential common regulators and mechanisms would be attractive therapeutic targets in these disorders.

Two of the cell types that undergo fusion, osteoclasts and multinucleated giant cells, are derived from a common progenitor from bone marrow and periphery. These cells are rendered fusion competent by common molecular mediators and ultimately mediate specialized functions in specific microenvironments. While OC can undergo fusion in both normal or pathological states such as Paget disease^32^, macrophages undergo fusion to form MGC primarily under inflammatory conditions such as chronic granulomatous disease^33^. However, the fact that the fusion of OC and MGCs are governed by common signaling mechanisms again makes these and their component molecules attractive targets for therapeutic intervention. Interestingly, we have previously shown that in response to ischemic injury, CD13^KO^ skeletal muscle satellite cells fused more readily than WT cells to form multinucleated myoblasts, suggesting that CD13 may also participate in fusion of other cell types, thus affecting processes such as skeletal muscle repair^16^.

Osteoclastogenesis comprises many steps from the commitment and survival of osteoclast progenitor cells, their differentiation into mononuclear pre-osteoclasts that fuse to generate multinucleated mature osteoclasts and finally activation of osteoclasts for bone resorption. Among the different steps, osteoclast fusion is thought to be the critical step in this phenomenon. Our data clearly indicate that osteoclast progenitor survival, differentiation and proliferation is not dependent on CD13 expression, suggesting that CD13 may be involved in the fusion mechanism to generate multinucleated osteoclasts.

The OC fusion process itself involves pre-fusion events; apposition of membranes via cell-adhesion molecules and the formation of an unstable intermediate stalk structure^34^. Fusion entails hemifusion of adjacent monolayers, remodeling of the phospholipid bilayer, mixing of membrane lipids, stalk elongation and formation of a bilayered diaphragm connecting the two cells. The post-fusion disruption of this diaphragm forms the fusion pore which expands in a dynamin-dependent manner producing a fused cell with continuous cytoplasm containing multiple nuclei. Studies have implicated various membrane-associated processes as critical to osteoclastogenesis such as clustering of membrane tetraspanins^35^ and endocytosis as well as specific molecules, such as DC-STAMP^27^, OC-STAMP^36^, dynamin and ATP6V0d2 (ATPase, H+ transporting V0 subunit d2)^37^. Recently, it has been shown that expression and localization membrane lipid species such as phosphatidylethanolamine^38^ and phophatidylserine^39^ are also essential for proper cell-cell fusion. We have shown that CD13 regulates dynamin-and clathrin-mediated endocytosis^21,22^, recycling of cell surface proteins^20^ and Src activation^40^, each of which have been shown to participate in OC fusion. Our data showing an aberrant persistence of fusion protein expression post-fusion in CD13^KO^ OCs suggests that CD13 may regulate the expression of these endocytic and fusion proteins during fusion, perhaps by mediating their internalization, endocytic trafficking, recycling and/or localization.

In addition, organizers of actin-based protrusions are pivotal in both osteoclast and macrophage fusion. Recently, we have reported that CD13 is a critical signaling platform that links the plasma membrane to dynamic mediators of actin cytoskeletal assembly and rearrangement^20^. It remains to be established if CD13 localization is required at the site of cell-cell fusion to enable cells to be fusion competent and that its expression on both cells is imperative for fusion.

In conclusion, in the present study we demonstrate that CD13 expression controls osteoclastogenesis specifically at the level of cell-cell fusion. Considering the diversity and importance of pathologies that are influenced by cell-cell fusion, identification of CD13-dependent molecular mechanisms and signaling that regulate myeloid fusion will provide novel therapeutic approaches in fusion pathologies.

## Legends

## Supplemental Data

**FIG S1. Flow sorting of primary murine bone marrow and spleen from WT and CD13**^**KO**^ **mice**. Live CD11b^lo^ CD115^hi^ Ly6G^+^ (CD3/B220/Nk1.1^)-^ cells were sorted to homogeneity using BD FACS ARIA and analyzed by FACS Diva. Data represents average of three independent experiments.

## Materials and Methods

### Animals

Global Wildtype and CD13^KO^ (C57BL/6J) male mice were generated and housed at the Gene Targeting and Transgenic Facility at University of Connecticut School of Medicine ^12^. All procedures were performed in accordance with the guidelines and regulations approved by the Institutional Animal Care and Use Committee.

### Reagents

Recombinant mouse M-CSF, RANKL, IL-4 and IL-13 were purchased from R&D Systems. Acid Phosphatase, Leukocyte (TRAP) Kit was purchased from Sigma-Aldrich. Giemsa/ MAY- GRÜNWALD Stain solution was purchased from Sigma-Aldrich. Toluidine Blue was obtained from EMD Millipore. Bovine cortical bone slices were a kind gift from Dr. Joseph Lorenzo, University of Connecticut School of Medicine^41^. Osteoplates were purchased from Corning.

### Antibodies

Antibodies to DC-STAMP1 (DCST1; Biorbyt, orb2242, rabbit polyclonal Ab), Dynamin (Abcam, ab3457, rabbit polyclonal Ab), Phalloidin-TRITC (Sigma-Aldrich, P1951), CD13 (SL13, Millipore, MABC950, rat monoclonal Ab)

### Flow cytometry

Antibodies for phenotypic analyses and sorting by flow cytometry used are as follows^24^; anti-CD3 (145-2C11), anti-B220 (RA3-6B2), anti-NK1.1 (PK136), anti-CD11b)M1/70), anti-CD115 (AFS98), anti-Ly6C (Al-21). All antibodies were purchased from Biolegend, BD Biosciences, e- Biosciences. UV Blue Live dead dye was purchased from Life Technologies. Labeling of cells for flow cytometry and sorting was performed as described. Briefly, flow cytometry on live cells were performed using BD-FACS Aria (BD Biosciences) and data analyzed with FlowJo software.

### In vivo analysis of WT and CD13^**KO**^ mice

### Micro-Computed Tomography and Histomorphometry^**41,42**^

Cortical and trabecular bone from 8-10 wks. old WT and CD13^KO^ mice male were isolated and scanned using micro-computed tomography (μCT) system and 3-D analysis and reconstruction were performed as described to measure the trabecular bone volume (BV/TV, %). For static histomorphometry, trabecular volume was measured as described. Osteoclasts were identified by multi-nucleated TRAP+ cells adjacent to the bone surface.

### Alkaline Phosphatase assay^**42**^

Osteoblast culture generated from bone marrow were stained for the alkaline phosphatase activity as described.

### Isolation of hematopoietic progenitor population from bone marrow and spleen

Bone marrow cells were obtained by flushing femur and tibia from WT or CD13^KO^ mice with 10ml 1xPBS and 2% heat inactivated FBS, followed by RBC lysis and filtering through 40um cell strainer (BD Biosciences). Total live cells counted with Countess Automated cell counter (Thermo Fisher Scientific) were stained with antibody cocktail at 4 degree C. Cells from mouse spleen was obtained by gentle crushing the organ between frosted microscopic slides in cold 10ml 1x PBS and 2% heat inactivated FBS.

### Generation and Culture of Osteoclast progenitors from bone marrow or spleen

Cells from mouse BM or spleen were stained with Ab cocktail containing anti-(CD3, B220, NK1.1, CD115, Ly6C) Ab and subjected to single-cell sorting by BD FACS Aria^24^. Sorted progenitors at a density of 10,000-50,000 cells/well were seeded in 96-well dish in α−MEM containing 10%FBS, 1% Penicillin-Streptomycin, 30ng/ml M-CSF and 30ng/ml RANKL at 37 degree C with 5% CO_2_ for 0-10d. Multi-nucleated osteoclasts were stained with Tartrate-Resistant Acid Phosphatase staining and assessed by counting cells with more than three nuclei. Average area of osteoclast was measured by ImageJ software.

### TRAP staining^**42**^

Osteoclast progenitor cells in α−MEM (GIBCO BRL) containing 10%FBS, 1% Penicillin-Streptomycin, 30ng/ml M-CSF and 30ng/ml RANKL grown on plastic or UV-sterilized, devitalized bovine cortical bone slices (placed in 96-well dishes), at a density of 50,000 cells/well for indicated time were fixed in 2.5% Glutaraldehyde and Tartrate-Resistant Acid Phosphatase in osteoclasts were stained using TRAP staining kit according to manufacturer’s instruction (Sigma-Aldrich).

### Bone resorption assay^**42**^

Osteoclast progenitors were seeded on Osteo Assay plate (Corning) at a density of 50,000 cells/well in α−MEM containing 10%FBS, 1% Penicillin-Streptomycin, 30ng/ml M-CSF and 30ng/ml RANKL for d10. Surface pit formation was measured by removing cells with 100ul of 10% bleach solution at RT for 5 min. Wells were washed with deionized water and allowed to dry. Cluster of pits formed was imaged using a light microscope (Olympus Scientific) and the area of resorption was measured by ImageJ software.

### Generation of Multinucleated Giant Cells (MGC) and Giemsa/ MAY-GRÜNWALD Staining

BM sorted Osteoclast progenitors were seeded on 96-well dishes at a density of 50,000 cells/well in α-MEM containing 10%FBS, 30ng/ml M-CSF for d5 followed by addition of and 30ng/ml IL-4 and 30ng/ml IL-13 for an additional d3. Cells were washed and fixed with 100% methanol followed by staining with Giemsa stain according to manufacturer’s instruction (Sigma-Aldrich).

### Immunofluorescence and microscopy

Osteoclast progenitors or MGC grown on glass coverslips that were previously coated with 5μg/ml fibronectin for indicated time period. Cells were fixed in 4% paraformaldehyde (Electron Microscopy Sciences) at RT for 30 min, permeabilized with 0.1% Triton-X-100 in PBS at RT for 5 min. Cells were blocked with blocking buffer containing 5% goat or donkey serum/5% BSA/1xPBS at RT for 1h followed by incubation with primary Ab in blocking buffer at 4 degree C for overnight. Cells were washed and treated with secondary Ab (1:1200) and DAPI (nuclear stain) in blocking buffer at RT for 1h. Coverslips were mounted with ProLong Gold antifade mounting medium (Life Technologies), visualized at excitation wavelength of 488nm (Alexa 488), 543nm (Alexa 594 or TRITC) and 405nm (DAPI) and imaged by Zeiss LSM 880 confocal fluorescence microscope.

### Immunoblot analysis

Cell lysates from flow-sorted bone marrow osteoclast progenitors grown in presence of M-CSF and RANKL over time were harvested in 1% NP40 lysis buffer containing 1X cOmplete Protease Inhibitor cocktail (Roche). Samples were separated by SDS-PAGE and transferred to nitrocellulose membrane, blocked in 1XTBST containing 5% bovine serum albumin, treated with primary Ab followed by appropriate secondary Ab and imaged by ChemiDoc Imaging system (Biorad). β actin was used as loading control.

### Statistical analysis

Statistical analysis was performed using unpaired, two-tailed Student’s t test using GraphPad Prism software and results are representative of mean ± SD. Differences at p≤ 0.05 were considered significant.

### Disclosures

All animal experiments in this study were reviewed and approved by Animal Care Committee at University of Connecticut Medical School. All the authors do not have any conflict of interest.

## Author contributions

MG, IK, HLA and LHS designed the research plan. MG, IK and HLA conducted experiments and analyzed the data. MG, HLA and LHS wrote the manuscript.

## Acknowledgments

We thank Dr. Joseph Lorenzo for the cortical bone slices and for the use of his light microscope. We also thank Susan Staurovsky and Evan Jellison from CCAM and Flow cytometry core facilities respectively at UConn Health, for providing technical assistance. This work was supported by National Institutes of Health grants R01HL127449 and R01HL125186 (to LHS and MG)

