## Supplemental File 1 for "CD13 is a Critical Regulator of Cell-cell Fusion in Osteoclastogenesis"

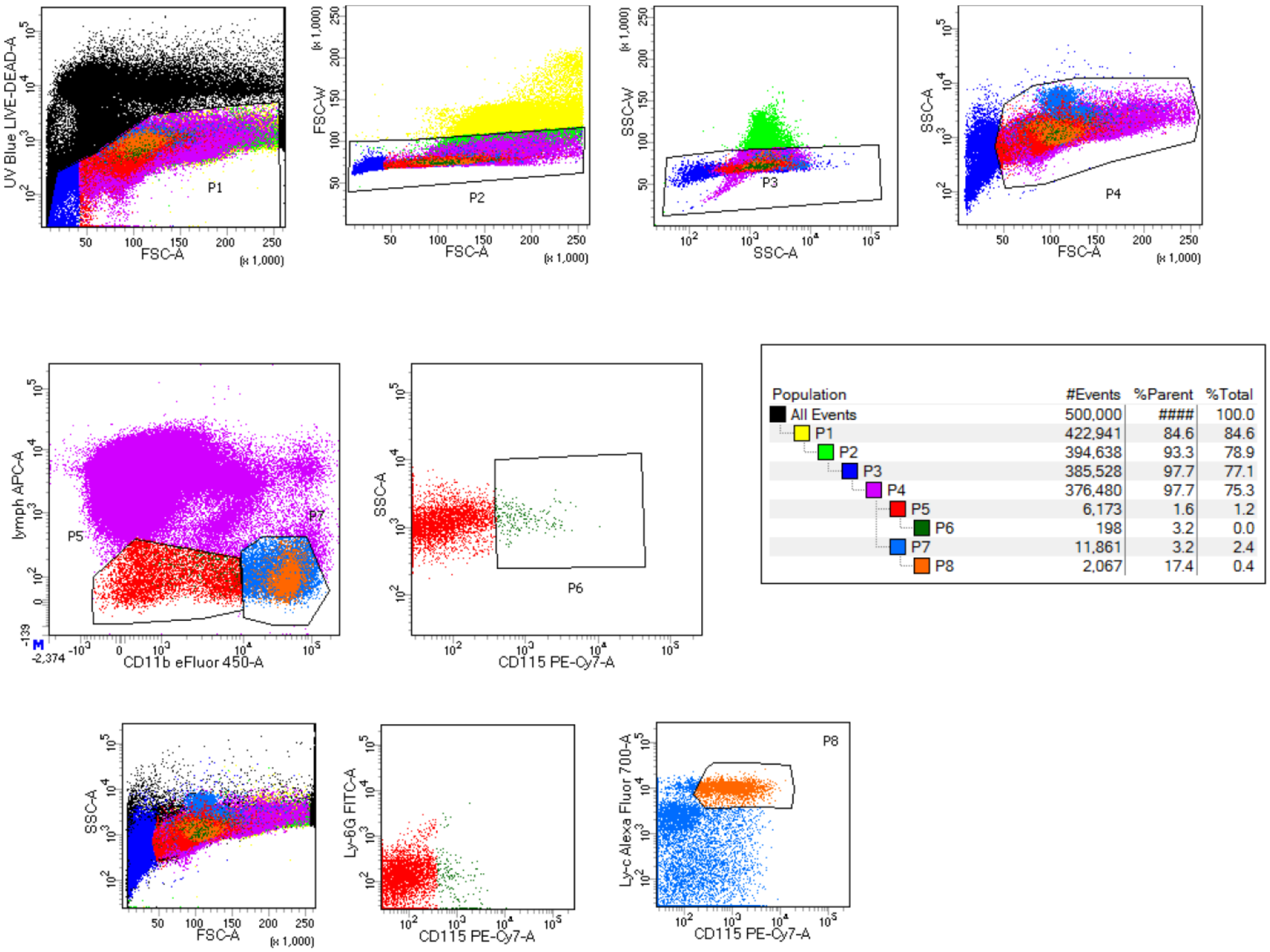


| Population | #Events | %Parent | %Total |
|---|---|---|---|
| All Events | 500,000 | #### | 100.0 |
| P1 | 422,941 | 84.6 | 84.6 |
| P2 | 394,638 | 93.3 | 78.9 |
| P3 | 385,528 | 97.7 | 77.1 |
| P4 | 376,480 | 97.7 | 75.3 |
| P5 | 6,173 | 1.6 | 1.2 |
| P6 | 198 | 3.2 | 0.0 |
| P7 | 11,861 | 3.2 | 2.4 |
| P8 | 2,067 | 17.4 | 0.4 |

**FIG S1. Representative image of flow sorting strategy of isolation of osteoclast progenitors from primary murine bone marrow or spleen from WT and CD13$^{KO}$ mice.** Live CD11b$^{lo}$ CD115$^{hi}$ Ly6G$^{+}$ (CD3/B220/Nk1.1$^{)-}$ cells were sorted to homogeneity using BD FACS ARIA and analyzed by FACSDiva.
